# Three separate pancreatic acinar cell receptors employ different combinations of intracellular pathways to Ca^2+^ signaling

**DOI:** 10.64898/2026.07.28.741212

**Authors:** Muhanad Salih, Julia V Gerasimenko, Oleg V Gerasimenko, Ole H Petersen

## Abstract

Repetitive cytosolic Ca2+ spikes in pancreatic acinar cells, elicited by low (physiological) concentrations of acetylcholine (ACh), cholecystokinin (CCK) and gastrin releasing peptide (GRP), control secretion of digestive enzymes, whereas high-intensity stimulation induces sustained Ca2+ elevation initiating acute pancreatitis. Since the discovery of the Ca2+-releasing function of inositol trisphosphate (IP_3_), it has generally been assumed that Ca2+ signaling relies on the phospholipase C – IP3 pathway. We have now compared the mechanisms of action of the three physiological stimulants, all acting on different receptors, but each coupled to the IP3 pathway. Unlike ACh, low physiological concentrations of CCK and GRP cannot elicit Ca2+ signals without co-operation of an additional intracellular mechanism. CCK-elicited Ca2+ signalling requires activation of intracellular receptors for nicotinic acid adenine dinucleotide phosphate (NAADP), whereas this is not the case for the action of GRP that nevertheless relies on the operation of CD38, the enzyme involved in the synthesis of both cyclic ADP ribose and NAADP. Even Ca2+ signals elicited by ACh are partially dependent on CD38. It is engagement of these additional non- IP3 pathways that allows low concentrations of secretagogues to elicit safe Ca2+ spiking and therefore secretion, obviating the need for potentially toxic high levels of agonists.

**Graphical abstract:** 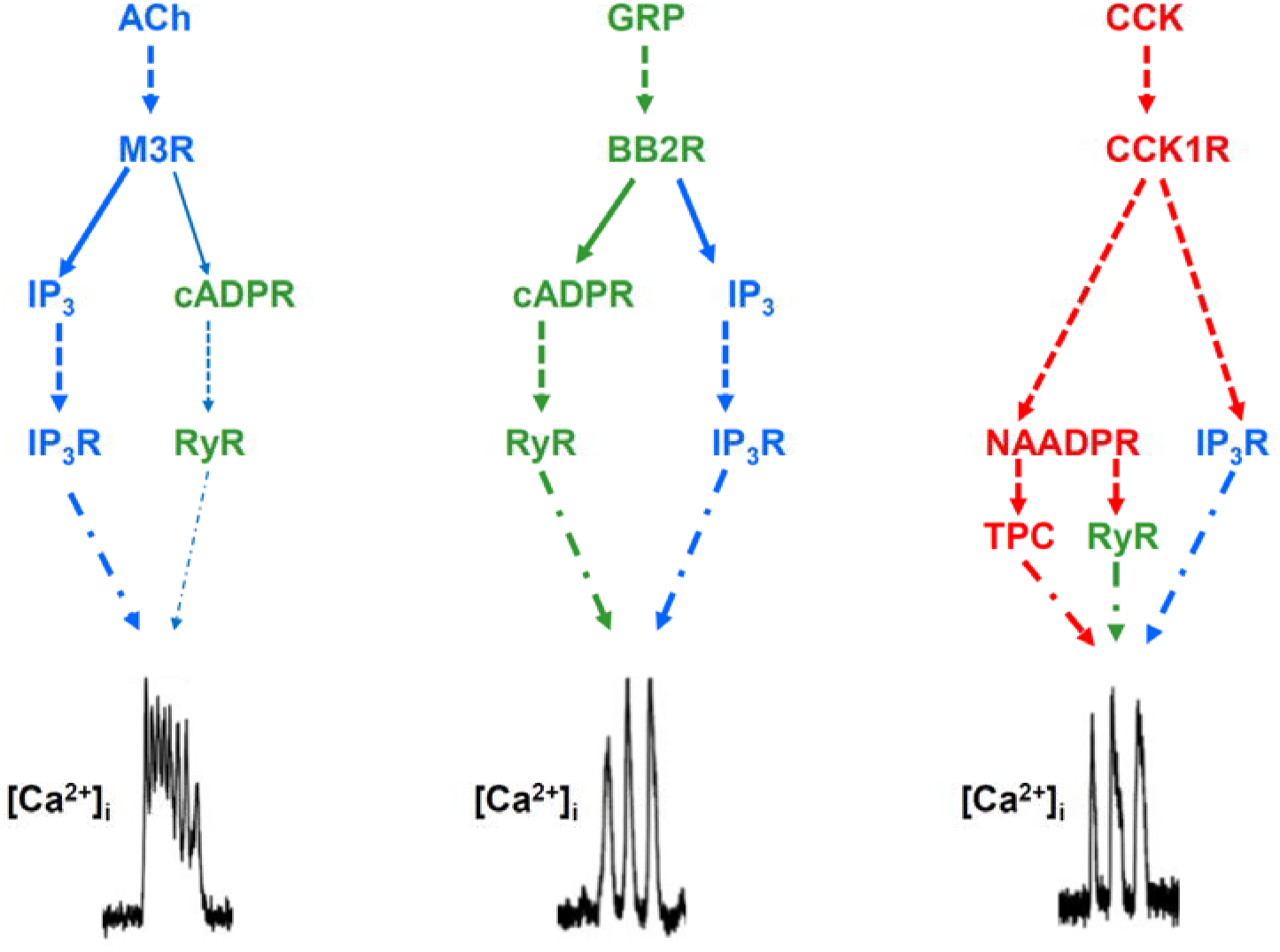

**In brief:** Three different agonists acting on three different receptors on the same cell employ different combinations of intracellular pathways to generate cytosolic Ca^2+^ signals

**Highlights:**

*At least two separate intracellular pathways and intracellular receptors are required for physiological Ca^2+^ signaling in normal pancreatic acinar cells
*Although functional IP_3_ receptors are always required, the second parallel pathway differs for the different agonists
*The exceptional sensitivity of pancreatic acinar cells to physiological, low pico-molar, CCK concentrations is due to sensitization of IP_3_ and NAADP receptors to resting messenger levels

## Introduction

Acetylcholine (ACh), cholecystokinin (CCK) and gastrin releasing peptide (GRP) are hormones and neurotransmitters with multiple functions in a multitude of peripheral organs as well as in the brain and spinal cord (1–5). The mechanisms of action of ACh and CCK has been intensively studied in the pancreatic acinar cell, which has long served as a key cell biological model for pioneering studies of protein synthesis, processing and exocytosis (6), epithelial ion channels (7) and Ca^2+^ signalling in electrically non-excitable cells (8). The physiological roles of ACh and CCK in digestion are well established. These agents elicit secretion of an enzyme-rich fluid from the pancreas in response to food intake. There is general agreement that ACh and CCK act on pancreatic acinar cells by interacting with muscarinic M3 and high-affinity CCK1 receptors, respectively, causing primary release of Ca^2+^ from intracellular stores, followed by opening of Ca^2+^ Release Activated Ca^2+^ (CRAC) channels in the plasma membrane, both processes serving to elevate the cytosolic Ca^2+^ concentration ([Ca^2+^]_i_) (9,10). It is the rise of [Ca^2+^]_i_ that activates exocytosis and opens ion channels important for fluid secretion (9,10). Whereas the physiological Ca^2+^ signals controlling secretion occur as repetitive Ca^2+^ spikes, sustained elevations of [Ca^2+^]_i_ are toxic, inducing cell damage leading to acute pancreatitis (9–11). GRP, and its frog skin analogue bombesin, also activate pancreatic enzyme and fluid secretion (12–14), interacting with receptor sites separate and different from those for CCK (15). There are several GRP receptor subtypes. It is the bombesin receptor subtype II (16) that mediates the physiologically relevant action of GRP. This receptor is overexpressed in many types of cancer, including prostate cancer, breast cancer, pancreatic cancer and lung cancer and is related to the growth of these malignancies (16–18).

Early studies of the mechanism by which CCK elicits cytosolic Ca^2+^ signals indicated that CCK used the same intracellular pathway as acetylcholine (ACh), namely activation of phospholipase C (PLC), resulting in generation of inositol 1,4,5- trisphosphate (IP_3_). This messenger would then bind to Ca^2+^-conducting IP_3_ receptors (IP_3_Rs) in the endoplasmic reticulum (ER) through which Ca^2+^ would be liberated from the store (19–24). Since then, it has been repeatedly confirmed that both ACh- and CCK-elicited Ca^2+^ signal generation depends completely on functional intracellular IP_3_Rs, but also on functional ryanodine receptors (RyRs) (9,10). However, another Ca^2+^-releasing messenger, nicotinic acid adenine dinucleotide phosphate (NAADP) (25), unknown at the time when the Ca^2+^ releasing function of IP_3_ was discovered (26), later turned out to play a critical role for the action of CCK but not for ACh (27–30). A third intracellular Ca^2+^ releasing messenger, cyclic ADP-ribose (cADPR), may also play a role. At first, it appeared that the action of CCK was blocked by a cADPR antagonist, 8-NH_2_-cADPR, that did not affect Ca^2+^ signal generation by ACh, but subsequent work indicated that the cADPR antagonist only inhibited CCK-elicited Ca^2+^ signalling in the absence of intracellular glucose (31). Further studies showed that ACh, like CCK, elicited cADPR production (30) and played a role in ACh-induced Ca^2+^ signal generation (32).

Although there are still many uncertainties about the intracellular steps activated by CCK (15), it is at least clear that the action of this hormone, at physiological concentrations (1-10 pM – refs 2,33), is completely dependent on both functional IP_3_ and NAADP receptors (15). In contrast, we know little about the mechanism of action of GRP. Here, we compare the effects of GRP with those of ACh and CCK on the cytosolic Ca^2+^ concentration ([Ca^2+^]_i_) in normal acutely isolated intact mouse pancreatic acinar cells, probing the differential effects of selective cell-permeant pharmacological agents. In spite of the very similar Ca^2+^ signal patterns evoked by CCK and GRP, it turns out that the action of GRP does not depend on functional NAADP receptors, but – like CCK and ACh – requires functional IP_3_Rs. We have investigated the effect of specifically inhibiting CD38, an enzyme involved in the synthesis of NAADP and cADPR (34). This consistently blocked GRP-elicited Ca^2+^ signal generation, whereas CCK could still evoke normal Ca^2+^ signals. These results indicate that although cytosolic Ca^2+^ signal generation elicited by both CCK and GRP require functional IP_3_Rs, the PLC-IP_3_ pathway alone is insufficient. For both the actions of CCK and GRP, an additional pathway needs to be recruited, but this additional pathway is different for these two agonists. We also show that the requirement for functional NAADP receptors and CD38 activity can be overcome at high agonist concentrations, outside the physiological range for peripheral hormone action (33), but relevant for neurotransmitter actions in the central nervous system (35,36). The engagement of Ca^2+^ releasing mechanisms additional to the established IP_3_ pathway enables Ca^2+^ signal generation, and therefore secretion, at remarkably low agonist concentrations. This is critically important because the high concentrations of CCK employed in the nervous system have been known for a long time to elicit toxic effects in pancreatic acinar cells (37,38). Ensuring safe Ca^2+^ signalling events in the pancreatic acinar cells therefore requires sensitivity to very low concentrations of stimulating agonists and our new results explain how this is achieved.

## Results

### GRP induces Ca^2+^ mobilization from intracellular stores in pancreatic acinar cells

Because of the absence of any detailed prior studies of GRP-elicited Ca^2+^ signalling, we started out by establishing the concentrations of GRP required to elicit cytosolic Ca^2+^ signals. As seen in Fig. 1A, 10 pM GRP had no effect, but a small and delayed effect of GRP was observed at 100 pM. At a concentration of 500 pM, the effect of GRP was close to maximal, although slightly broader Ca^2+^ spikes could be seen at the higher GRP concentrations of 1 and 10 nM (Fig. 1A). The pattern and amplitude of the Ca^2+^ signals elicited by 125 pM GRP were comparable to those elicited by 5 pM CCK (Fig. 1B). It is well established that stimulant-elicited Ca^2+^ signal initiation in pancreatic acinar cells does not depend on the presence of external Ca^2+^, as the signals are the result of release of Ca^2+^ from internal stores (9). However, every Ca^2+^ spike causes some extrusion of Ca^2+^ (39), mediated by plasma membrane Ca^2+^ pumps (40), and therefore Ca^2+^ entry is needed in the long term to compensate for this loss (9). Eventually, continuous Ca^2+^ spiking elicited by sustained stimulation becomes dependent on external Ca^2+^, but this can take a long time (39), particularly at low agonist concentrations, due to reuptake of Ca^2+^ into the ER (9,40) during inter-spike intervals. In our experiments, GRP-elicited Ca^2+^ signalling did not require the presence of Ca^2+^ in the external solution within the time frame of our recordings (Fig. 1C, 7/7 cells) and this was also the case for the action of CCK (Fig. 1D, 6/6 cells).

**Figure 1.**
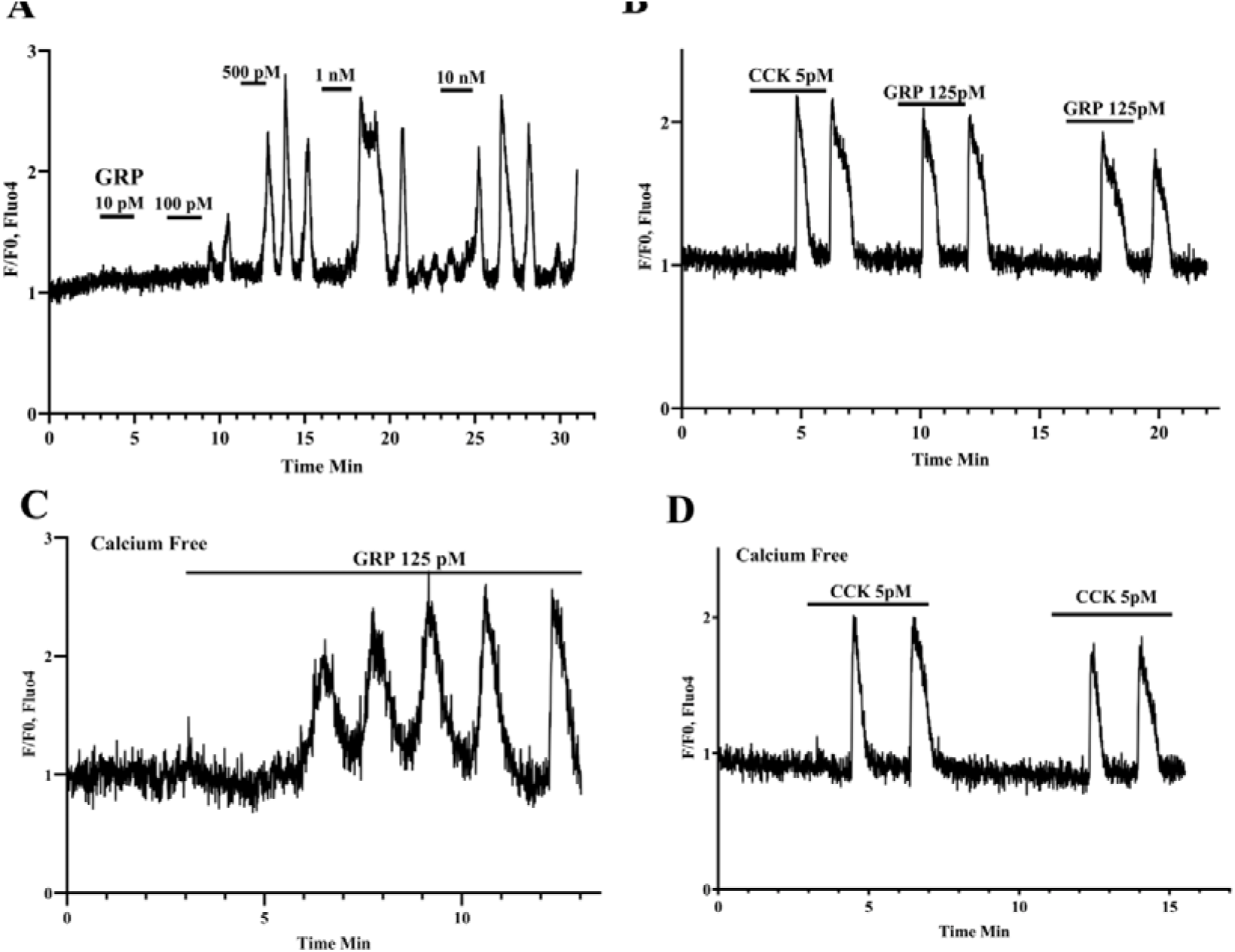
GRP-induced Ca²⁺ spiking in mouse pancreatic acinar cells. **(A)** GRP, at a concentration of 10 pM, failed to elicit Ca²⁺ signals, but evoked small, delayed Ca²⁺ spikes at 100 pM. Increasing the GRP concentration to 500 pM markedly increased the spike amplitudes. Higher concentrations, 1 and10 nM, increased spike durations without altering spike amplitude. **(B)** The Ca²⁺ signals induced by 125 pM GRP closely resembled those evoked by 5 pM CCK, showing similar oscillatory patterns and amplitudes. **(C, D)** Both GRP and CCK initiated Ca²⁺ signalling in the absence of extracellular Ca²⁺. Intracellular Ca²⁺ signals were monitored using Fluo-4 fluorescence and are presented as normalised fluorescence ratios (F/F₀).

### GRP-elicited Ca^2+^ signals depend on functional IP_3_ receptors and G_q/11_

It is well-established that CCK- and ACh- elicited Ca^2+^ signaling can be blocked by caffeine, acting as a membrane-permeant inhibitor of IP_3_Rs (9,23,41). Ca^2+^ signal generation elicited by both 500 pM GRP (Fig. 2A) and 1 nM GRP (24/24 cells) was abolished by 20 mM caffeine. This inhibition was fully reversible (Fig. 2A), consistent with previous data on the actions of ACh and CCK (9,10). CCK- elicited Ca^2+^ signal generation can be mimicked by GTP-γ-S (42) and abolished by YM-254890, a highly selective inhibitor of G_q/11_ (43). YM-254890 is a natural product isolated from *Chromobacterium* sp., that was initially identified as a potent inhibitor of ADP-induced human platelet aggregation through disruption of P2Y₁-mediated receptor signalling (44,45) and later characterised as a specific inhibitor of G_q/11_ (43). We tested the effect of this inhibitor on GRP-elicited Ca^2+^ signaling and showed that YM-254890 (1 μM) abolished Ca^2+^ signal generation (Fig. 2B). Stimulation with 125 pM GRP produced robust Ca²⁺ signals, which were abolished by 1 µM YM-254890 in all tested cells (Fig. 2B). Similar results were obtained at the higher GRP concentration of 500 pM (Fig. 2B). The inhibitory effect of YM-254890 was irreversible as Ca²⁺ oscillations did not recover following washout.

**Figure 2.**
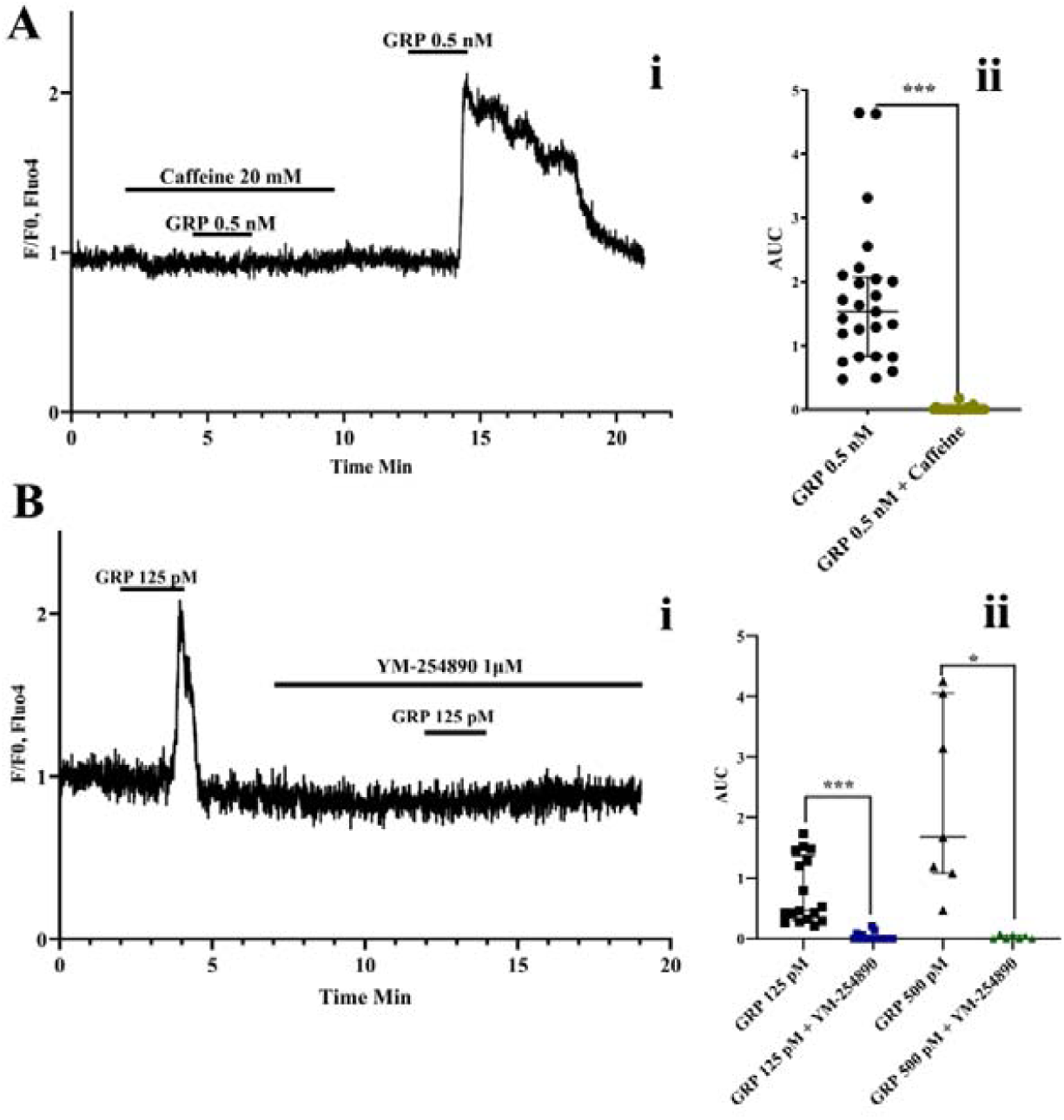
Requirement for G_q/11_ activation and functional IP₃ receptors in GRP-evoked Ca²⁺ signaling. **(Ai)** Ca²⁺ spikes evoked by 500 pM GRP were reversibly inhibited by caffeine (20 mM). **(Aii)** Quantification of Ca²⁺ signals before, and after caffeine application (25 cells). **(Bi)** inhibition of G_q/11_ activation by 1 μM YM-254890 abolished Ca²⁺ mobilisation elicited by 125 pM GRP. **(Bii)** Quantification of GRP-evoked Ca²⁺ responses in the absence and presence of YM-254890. Ca²⁺ mobilisation induced by 125 pM GRP was abolished in 17/17 cells and by 500 pM GRP in 8/8 cells. Intracellular Ca²⁺ signals are expressed as normalised fluorescence ratios (F/F₀). Data are shown as median ± IQR. Normality was assessed using the Shapiro–Wilk test, and statistical comparisons were performed using the Wilcoxon matched-pairs signed-rank test. *P < 0.05; ***P < 0.001.

### Ca^2+^ spiking evoked by GRP does not depend on functional NAADP receptors

As shown in Fig 1B, CCK (5 pM) and GRP (125 pM) evoke very similar patterns of cytosolic Ca²⁺ spiking. We therefore wanted to test whether these two agents also employ the same intracellular messenger pathways. In the case of CCK, it has been shown that blockade of NAADP receptors by the highly selective inhibitor Ned-19 (46) abolished Ca²⁺ signal generation, whereas it had no effect on signal generation by agents acting on muscarinic receptors (47,48). First, we re-examined the effect of Ned-19 on CCK-induced Ca²⁺ signalling. At a CCK concentration of 5 pM, Ca²⁺ spiking was abolished by Ned-19. Increasing the CCK concentration to 50 pM resulted in small residual spikes, whereas further elevation of the CCK concentration to 500 pM led to a sustained [Ca²⁺]_i_ elevation with super-imposed spiking in the continued presence of Ned-19 (Fig. 3A). Although this has not been shown before, this result was anticipated because the high concentration of CCK (500 pM), in contrast to the physiologically relevant concentrations (1-10 pM), results in IP_3_ production (49). At a high, un-physiological, CCK concentration, the level of intracellular IP_3_ is evidently sufficient to generate Ca²⁺ signals without recruitment of functional NAADP receptors.

**Figure 3.**
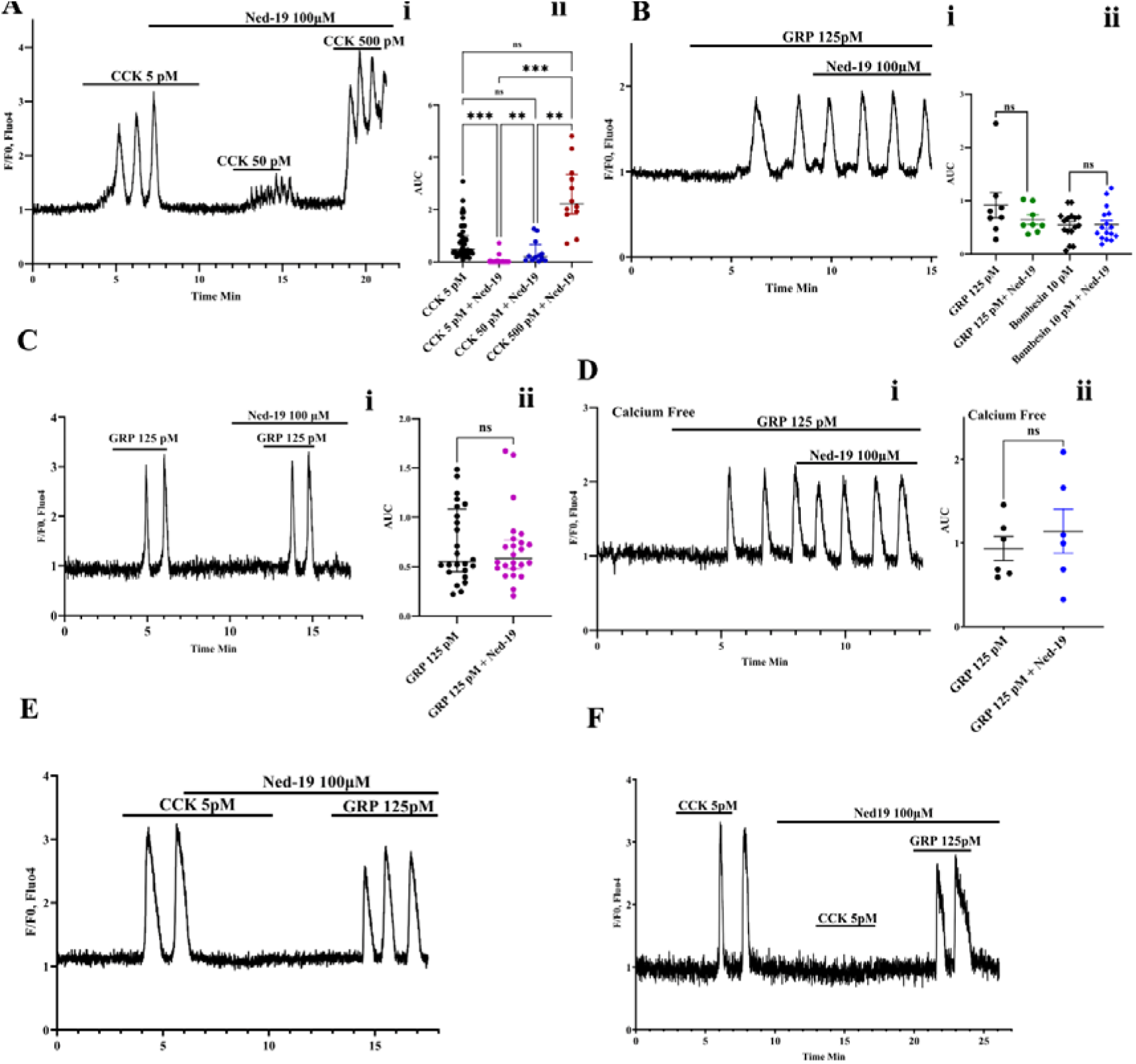
CCK, but not GRP, requires functional NAADP receptors for generation of Ca²⁺ spikes, whereas NAADP receptor inhibition enhances ACh-evoked Ca²⁺ signaling. **(Ai)** Ned-19 abolished Ca²⁺ spiking evoked by 5 pM CCK, but this inhibition was overcome by increasing the CCK concentration, first to 50 and then 500 pM in the continued presence of the inhibitor. **(Aii)** Quantification of CCK-evoked Ca²⁺ signals under the indicated conditions. **(Bi)** Ned-19 did not affect Ca²⁺ spiking elicited by 125 pM GRP in presence of extracellular calcium. **(Bii)** No difference in Ca²⁺ signaling before and after Ned-19 application, elicited by GRP (8/8 cells) or bombesin (10 pM) (16/16 cells). **(Ci)** In the presence of Ned-19, GRP evoked normal Ca²⁺ spiking. **(Cii)** No significant difference in Ca²⁺ spiking was observed between stimulation with GRP alone and in the presence of Ned-19**. (Di)** Ned-19 did not affect Ca²⁺ responses induced by 125 pM GRP in absence of extracellular calcium. **(Dii)** No difference in GRP-induced Ca²⁺ signaling before and after Ned-19 application in absence of extracellular Ca²⁺ (6/6 cells). **(E)** Ned-19 abolished ongoing CCK-elicited Ca²⁺ spiking, whereas GRP was able to evoke repetitive Ca²⁺ spiking in the same cell in the continued presence of Ned-19 (18/18 cells). **(F)** Ned-19 prevented the initiation of CCK-elicited Ca²⁺ spiking (29 of 34 cells; in 5 cells markedly reduced responses), while GRP evoked normal repetitive Ca²⁺ spiking inthe same cell in the continued presence of Ned-19. Intracellular Ca²⁺ signals were monitored using Fluo-4 fluorescence and are expressed as normalised fluorescence ratios (F/F₀). Data are presented as Median ± IQR except for (Dii) where presented as mean ± SEM. Statistical analyses were performed using the Kruskal–Wallis test with Dunn’s multiple-comparisons test for (Aii), paired *t*-test for (Dii), and Wilcoxon matched-pairs signed-rank test for the rest of comparisons, following assessment of normality with the Shapiro–Wilk test. \**P* < 0.05, **P < 0.01, \*\*\**P* < 0.001.

In contrast to the abolition of CCK-evoked Ca²⁺ signalling by Ned-19, GRP (125 pM)- induced [Ca²⁺] spiking remained unchanged in the presence of Ned-19 (100 µM), whether Ned-19 was added during ongoing oscillations (Fig. 3B) or applied before GRP stimulation (Figure 3C). Ned-19 also failed to interrupt or diminish Ca²⁺ signaling in the absence of external Ca²⁺ (Fig. 3D). The inability of Ned-19 to interfere with GRP-elicited Ca²⁺ signalling was confirmed by direct comparisons with the action of CCK in the same cell. CCK (5 pM)–induced oscillations were acutely interrupted by Ned-19, while GRP (125 pM) added subsequently elicited a normal Ca²⁺ signalling response (Fig. 3E). This difference between the effects of Ned-19 on the actions of CCK and GRP was further supported by a slightly different experimental protocol in which Ned-19 (100 µM) abolished the ability of CCK to initiate Ca²⁺ spiking while GRP could still evoke a large response (Fig. 3F). We also checked the effect of Ned19 on bombesin (10 pM)-induced Ca²⁺ oscillations. Consistent with expectations, Ned-19 did not alter the Ca²⁺ spiking response evoked by 10 pM bombesin (Fig. 3Bii). Fig. 3 shows that there is a very clear difference between the effect of Ned-19 on CCK- and GRP (bombesin)-elicited Ca²⁺ signal generation, since the physiological action of CCK has an absolute requirement for functional NAADP receptors whereas GRP elicits normal Ca²⁺ spiking also in the case of full NAADP receptor blockade.

### Ca^2+^ signalling elicited by CCK, GRP, and ACh showed different levels of dependency on CD38

Despite the possible involvement of various enzymes in cADPR and NAADP synthesis, CD38 remains the only enzyme characterised to mediate the production of both these messengers (34,50,51). 78c (CD38-IN-78c) is a selective, reversible, non-competitive inhibitor of murine and human CD38 that inhibits both its hydrolase and cyclase activities (52–54). Ca²⁺ oscillations triggered by CCK (5 pM) were not affected by 30 μM 78c (Fig. 4A), and remained unaffected even when the concentration of 78c was increased to 50 µM (26/26). In stark contrast, Ca²⁺ spiking induced by GRP (125 pM) was abolished by the same concentration of 78c (Fig. 4Bi), with full inhibition observed in all cells tested (Fig. 4Bii). This differential sensitivity was most convincingly demonstrated by direct comparisons in the same cell, where the GRP-induced response was blocked while CCK-induced oscillations were normal (Fig. 4C). The inhibitory effect of 78c on GRP-induced Ca^2+^ spiking could be overcome by increasing the concentration of GRP. Both 1.25 nM and 12.5 nM GRP were able to elicit Ca²⁺ signals in the presence of 30 µM 78c (Fig. 4D). As already mentioned, removal of extracellular Ca²⁺ had little effect on CCK-induced Ca²⁺ spiking (Fig 1D) but, under Ca²⁺-free conditions, application of 30 µM 78c almost completely blocked CCK-induced Ca²⁺ spiking in the majority of cells (Fig. 4E; 26/38). A smaller proportion of cells (10/38) showed a marked reduction in CCK-elicited spiking activity (Fig. 4E), while only 2 out of 38 cells had normal signals. Interestingly, quantification of the responses revealed that cells showing only partial inhibition by 78c exhibited significantly larger initial CCK-evoked responses than cells in which 78c produced complete inhibition (Fig. 4F). Highly CCK-responsive cells are less susceptible to full suppression by 78c.

**Figure 4.**
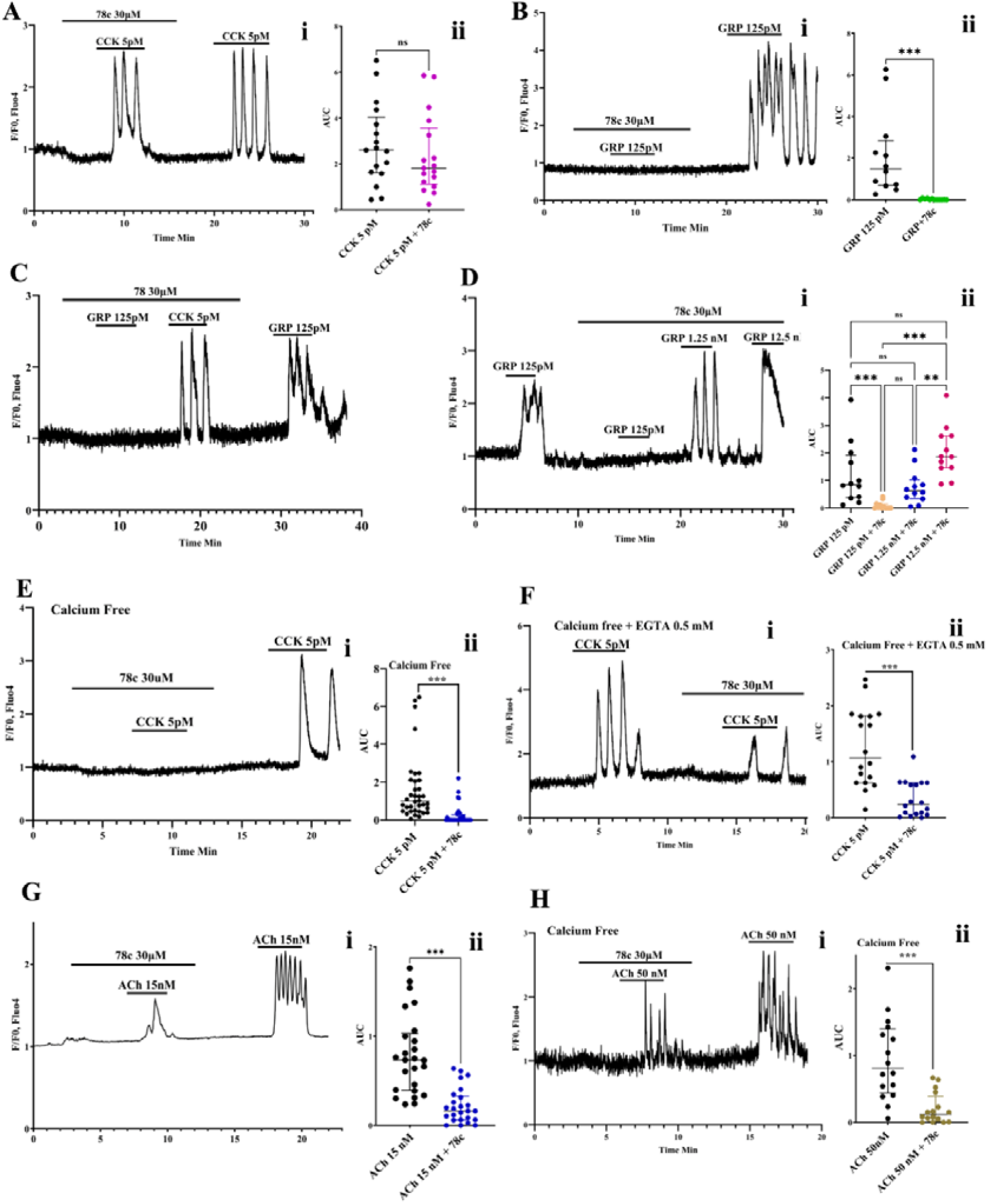
Differential dependence of CCK-, GRP-, and ACh-elicited Ca²⁺ signalling on CD38 activity. **(Ai)** Inhibition of CD38 with 30 μM 78c did not prevent normal Ca²⁺ spiking evoked by 5 pM CCK in the presence of extracellular Ca²⁺. **(Aii)** Quantification of CCK-evoked Ca²⁺ signaling under the indicated conditions. **(Bi)** In the presence of 30 μM 78c, Ca²⁺ spiking evoked by 125 pM GRP was abolished. **(Bii)** Quantification of GRP-evoked Ca²⁺ signaling with and without 78c. **(C)** In the same cell, the GRP-evoked response was abolished (25/25) whereas the CCK-induced spiking persisted in the continued presence of 78c. **(Di)** Increasing the GRP concentration restored Ca²⁺ spiking despite continued exposure to 78c. **(Dii)** Quantification of GRP-evoked Ca²⁺ responses at the indicated GRP concentrations in the absence and presence of 78c. **(Ei)** Removal of extracellular Ca²⁺ revealed a requirement for CD38 activity in CCK-elicited Ca²⁺ signalling, as 78c markedly reduced or abolished Ca²⁺ spiking. **(Eii)** Quantification of CCK-evoked Ca²⁺ signaling under Ca²⁺-free conditions with and without 78c. **(Fi)** The dependence of CCK- induced Ca²⁺ signalling on CD38 activity was confirmed in Ca²⁺-free external solution containing EGTA, where 78c markedly suppressed the response. **(Fii)** Quantification of CCK-elicited Ca²⁺ responses under the indicated conditions. **(Gi)** 78c reduced both the frequency and amplitude of ACh-induced Ca²⁺ spikes in the presence of extracellular Ca²⁺. **(Gii)** Quantification of ACh-evoked Ca²⁺ responses with and without 78c. **(Hi)** In the absence of extracellular Ca²⁺, 78c reduced ACh-induced Ca²⁺ spiking. **(Hii)** Quantification of ACh-evoked Ca²⁺ responses under Ca²⁺-free conditions with and without 78c. Intracellular Ca²⁺ signals were measured using Fluo-4 fluorescence and are expressed as normalised fluorescence ratios (F/F₀). Data are presented as median ± IQR. Statistical analyses were performed using the Friedman test with Dunn’s multiple-comparisons test for **(Dii)** and the Wilcoxon matched-pairs signed-rank test for all other comparisons, following assessment of normality with the Shapiro–Wilk test. \**P* < 0.05, **P < 0.01, \*\*\**P* < 0.001.

The same overall pattern was observed when 0.5 mM EGTA was added to the Ca²⁺- free external solution to chelate residual extracellular Ca²⁺. Under these conditions, CCK-induced Ca²⁺ oscillations were maintained. However, the response to CCK was abolished in 10 of the 22 cells by 30 µM 78c, while the remaining 12 cells exhibited a reduced response (Fig. 4F). The cells that remained partially responsive in the presence of 78c displayed significantly larger initial CCK-evoked responses than cells in which spiking was completely abolished (Fig. 4F).

Unexpectedly, 30 µM 78c markedly inhibited ACh-induced Ca²⁺ spiking at low agonist concentrations, with consistent reductions observed at both 20 nM ACh (25/28 cells - the remaining 3 cells did not show substantially reduced signals), and 15 nM ACh (Fig. 4G; 25/25 cells). In Ca²⁺-free conditions, pancreatic acinar cells exhibited reduced sensitivity to ACh, and only concentrations of 50 nM or higher elicited measurable Ca²⁺ oscillations. Despite the higher concentration of ACh required under these conditions, 78c produced a similar inhibitory effect (Fig. 4H). However, unlike in the presence of extracellular Ca²⁺, in which we only observed a reduction, 30 µM 78c - under Ca²⁺-free conditions - abolished ACh-induced Ca²⁺ oscillations in 8 of 16 cells, while the remaining cells displayed a reduction in amplitude and length of Ca²⁺ spikes (Fig. 4H).

### Ned-19 potentiates ACh-evoked Ca²***⁺*** signals

We confirmed that Ned-19 had little or no effect on ongoing ACh-elicited Ca²⁺ spiking when applied after the response had been initiated (Fig. 5A, 14/14 cells) (47). However, addition of Ned-19 (100 µM) before stimulation with ACh (30 nM) produced a markedly different outcome. Under these conditions, ACh-evoked Ca²⁺ responses were enhanced, with significant increases in both spike amplitude and spike frequency (Fig. 5B, 28/28 cells). To further investigate this sensitising effect, we examined whether Ned-19 could counteract the inhibitory actions of the CD38 inhibitor 78c. As already shown in Fig. 4G,H, 78c (30 μM) reduced both the amplitude and frequency of ACh-induced Ca²⁺ spikes. Fig. 5C (26/26 cells) shows that Ned-19 restored the normal Ca²⁺ signal generation that had been markedly reduced by 78c. Consistent with this finding, 78c also attenuated the enhancement of ACh-induced Ca²⁺ spiking produced by Ned-19 (Fig. 5D, 10/10 cells).

**Figure 5.**
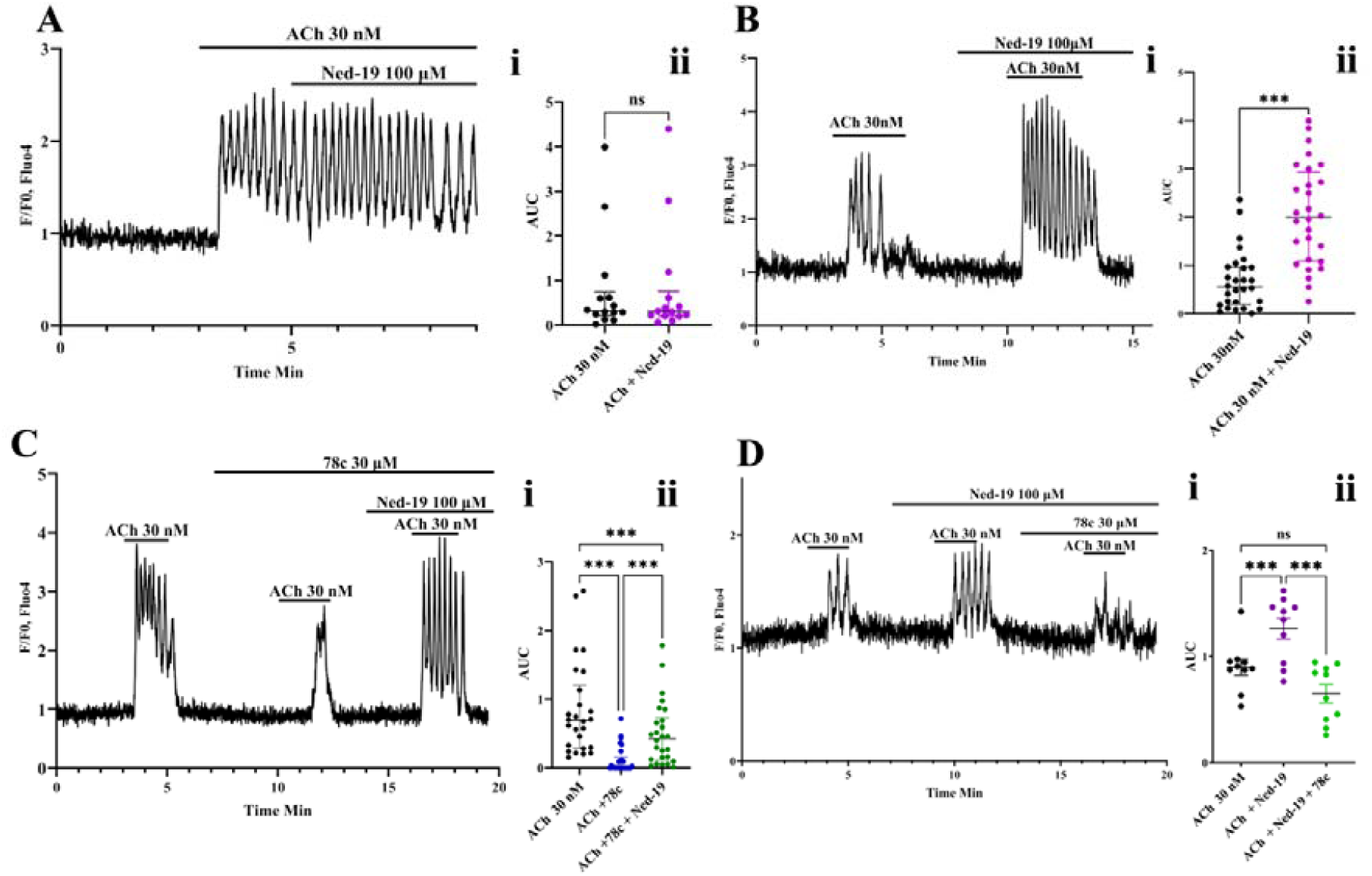
Ned-19 potentiates ACh-elicited Ca²⁺ mobilization. **(Ai)** Application of Ned-19 during ongoing ACh-evoked Ca²⁺ oscillations did not alter the frequency or amplitude of the Ca²⁺ spikes. **(Aii)** Quantification of Ca²⁺ signalling under the indicated conditions. **(Bi)** In contrast, addition of Ned-19 before stimulation with 30 nM ACh enhanced both the frequency and amplitude of Ca²⁺ spikes. **(Bii)** Quantitative analysis demonstrating a significant increase in ACh-evoked Ca²⁺ responses in the presence of Ned-19. **(Ci)** The reduction of ACh-elicited Ca²⁺ signals by 78c was reversed by Ned-19. **(Cii)** Quantification of ACh-elicited Ca²⁺ responses under the indicated conditions. **(Di)** 78c attenuated the potentiating effect of Ned-19 on ACh-induced Ca²⁺ responses. **(Dii)** Quantification of ACh-elicited Ca²⁺ responses under the indicated conditions. Intracellular Ca²⁺ signals were measured using Fluo- 4 fluorescence and are expressed as normalised fluorescence ratios (F/F₀). Data are presented as median ± IQR except (Dii) where presented as mean ± SEM. Statistical analyses were performed using the Wilcoxon matched-pairs signed-rank test for **(Aii)** and **(Bii)**, the Friedman test with Dunn’s multiple-comparisons post hoc test for **(Cii)**, and one-way ANOVA with Tukey’s multiple-comparisons test for **(Dii)** following assessment of normality using the Shapiro-Wilk test. \**P* < 0.05, \*\***P** < 0.01, \*\*\***P** < 0.001.

## Discussion

Our results show that the low (physiological) concentrations of secretagogues that need to be employed for safe Ca²⁺ signaling - repetitive Ca²⁺ spiking rather than toxic sustained [Ca²⁺]_i_ elevation (9,10) – require two different intracellular pathways to release Ca²⁺ from intracellular stores. Surprisingly, the two peptides, CCK and GRP, engage different pathways. Although it is often assumed that the PLC-IP_3_-IP_3_R pathway is responsible for the ability of CCK to elicit cytosolic Ca²⁺ signals in their target cells (2,24,43), it is already known that in the pancreatic acinar cells, the operation of the IP_3_ pathway alone is insufficient for CCK, at physiologically relevant concentrations (1-10 pM – ref. 33), to generate cytosolic Ca²⁺ signals. In addition to functional IP_3_Rs, it also requires functional NAADP receptors (9,10,27–30,34,47). Our new data confirm that functional NAADP receptors are required for CCK-elicited Ca²⁺ signalling (Fig. 3) but, unexpectedly, show that pharmacological inhibition of CD38, the enzyme involved in the production of both NAADP and cADPR (34), does not affect CCK-elicited Ca²⁺ signal generation, in the presence of a normal extracellular Ca²⁺ concentration (Fig. 4). This is surprising because it is known that knock-out of CD38 prevents CCK-elicited NAADP formation (34). Thus, it would appear that Ca²⁺ signalling evoked by physiological CCK concentrations depends on functional NAADP receptors but does not require CCK-elicited NAADP production. Likewise, CCK-elicited Ca²⁺ signalling depends on functional IP_3_Rs (43) but not on CCK-elicited IP_3_ production, because there is no rise in IP_3_ levels at CCK concentrations below 10 pM (49). Our new results (Fig. 2) show that cytosolic Ca²⁺ signal generation elicited by low concentrations of GRP depends on G_q/11_ activation and functional IP_3_Rs, as also shown for the action of CCK. However, in stark contrast to the situation for CCK, we now show that the action of GRP is completely dependent on the operation of CD38, but does not require functional NAADP receptors (Figs. 3 and 4), suggesting that cADPR and its receptor most likely are critical for the ability of GRP to elicit Ca²⁺ signals. Table 1 summarizes the effects of the four membrane-permeant inhibitors on the actions of CCK, GRP and ACh, respectively Our new result showing that CCK-elicited Ca²⁺ signalling works normally when CD38 is inhibited pharmacologically may seem surprising, but is in agreement with the finding of Cosker et al (34) that CCK-elicited Ca²⁺ signalling is normal in CD38^-/-^ acinar cells. What then is the trigger for the CCK-elicited release of Ca²⁺ from intracellular stores? One possibility could be cADPR, as physiological concentrations of CCK have been shown to increase the cADPR level in pancreatic acinar cells (30). This would fit in with early patch-clamp whole cell recording data showing that inhibition of intracellular cADPR receptors by the membrane impermeant agent 8- NH2-cADPR abolished CCK-elicited oscillations of Ca²⁺-dependent ion currents. However, it turned out that this only happens if there is no glucose in the intracellular solution. In the presence of glucose, 8-NH2-cADPR did not inhibit CCK-elicited spiking of Ca²⁺-activated ion currents (31). Given that CCK-elicited Ca²⁺ signalling is absolutely dependent on both functional IP_3_Rs and functional NAADP receptors, the most likely explanation is that factors sensitizing both IP_3_Rs and NAADP receptors to the resting levels of IP_3_ and NAADP are at work. Although both CCK1 and CCK2 receptors are coupled to G_q/11_ there is also evidence for coupling to many other G-proteins (2). Sensitization of IP_3_Rs to the resting level of IP_3_, resulting in Ca²⁺ signal generation without an increase in IP_3_ production, can be induced by the SH-group oxidising agent thimerosal (55,56) and this has also been shown to occur in normal pancreatic acinar cells, where thimerosal elicits large CCK-like cytosolic Ca²⁺ spikes that can be blocked by caffeine (57). Many different regulators of the sensitivity of IP_3_Rs to IP_3_ have been described (58) and further work is required to identify which ones may be involved in the action of CCK. In contrast to the wealth of information, at the molecular level, about the IP_3_Rs, that directly conduct Ca²⁺ (58), we know relatively little about the NAADP receptors. They are not Ca²⁺ conducting ion channels, but activate two-pore channels (TPC) in acid intracellular Ca²⁺ stores, including endosomes, lysosomes and secretory granules as well as ryanodine receptors in both the ER and acid stores (47,48,59,60).

**Table 1.** The effects of four different membrane permeant inhibitors on Ca^2+^ signals elicited by CCK (at physiologically relevant concentrations, 1-10 pM), GRP (low concentrations) or ACh (low concentrations)

|  | **Caffeine** | **YM-254890** | **Ned-19** | **78c** |
| --- | --- | --- | --- | --- |
| **CCK** | Blocked | Blocked | Blocked | Normal* |
| **GRP** | Blocked | Blocked | Normal | Blocked |
| **ACh** | Blocked<sup>1</sup> | Blocked<sup>2</sup> | Enhanced | Reduced |

\*But in the absence of extracellular $\text{Ca}^{2+}$ : blocked; <sup>1</sup>Ref 72; <sup>2</sup>Ref 43

In the absence of extracellular Ca²⁺, CCK-elicited Ca²⁺ signal generation, which was not inhibited by the Ca²⁺-free external solution itself, became dependent on CD38, as pharmacological inhibition of this enzyme, in this particular condition, abolished the ability of CCK to elicit Ca²⁺ spiking (Fig. 4). Although at first surprising, this is in agreement with the findings of Cosker et al (34), who showed that in acinar cells from CD38^-/-^ mice, CCK could not elicit Ca²⁺ spiking in the absence of external Ca²⁺, whereas this occurred normally in the presence of Ca²⁺. Thus, in the absence of external Ca²⁺, CD38 function becomes a necessity for signalling, implying a requirement for an increase in the NAADP concentration to trigger Ca²⁺ release. In the normal presence of external Ca²⁺, it would seem that a small CCK-elicited entry of Ca²⁺ (61) can compensate for the lack of CD38 activity. In this context, it is interesting that thimerosal-elicited Ca²⁺ signal generation in pancreatic acinar cells could not be induced in the absence of external Ca²⁺, but required a brief period of exposure to a normal external Ca²⁺ concentration for initiation of Ca²⁺ spiking (57). The nature of the Ca²⁺ influx pathway, that in these situations seems to be required to enable release of Ca²⁺ from intracellular stores, is unknown. In addition to the Orai1 type Ca²⁺ Release Activated Ca²⁺ (CRAC) channels (62), Piezo1 and TRPV4 channels (63,64) as well as non-selective cation channels (65) - most likely TRPM4 channels (66) - have been characterized in normal pancreatic acinar cells. One or more of these pathways could in principle be utilized, although it is unclear how these channels would be activated to mediate the Ca²⁺ influx that appears to be required in this situation. Endocytic Ca²⁺ uptake (67) may provide a more likely explanation and it has been proposed that endocytosis is of functional importance for CCK-elicited Ca²⁺ signal generation (68).

The mechanism of action of GRP is very different from that of CCK. The abolishment of GRP-elicited Ca²⁺ signalling when CD38 is inhibited by 78c (Fig. 4) can most simply be explained by a need for production of cADPR, as functional receptors for NAADP are not required (Fig. 3). This is in accordance with predictions from an earlier patch clamp study in which 8-NH2-cADPR was shown to abolish bombesin- elicited spiking of Ca²⁺-activated Cl^-^ current (69). Our finding that the inhibitory effect of 78c could be overcome by markedly increasing the GRP concentration could be due to a large increase in IP_3_ production at high GRP levels (70), obviating the need for recruiting cADPR to release Ca²⁺ from stores.

ACh action on muscarinic receptors provided the original, now classical, case for IP_3_- mediated Ca²⁺ signalling based on studies of pancreatic acinar cells (8,26). However, our new results showing that 78c reduces the amplitude of the Ca²⁺ spikes elicited by the muscarinic agonist, adds evidence to previous findings of Fukushi et al (32) and Yamasaki et al (30) indicating that there is also a role for cADPR. Our observation, that Ned-19 can markedly enhance ACh-elicited Ca²⁺ signals (Fig. 5), was entirely unexpected, but provides further evidence of linkages between the classical IP_3_ pathway and the more recently discovered additional mechanisms for Ca²⁺ signal generation. It may signify that if the NAADP pathway were to be blocked for some reason, then the cells can compensate by boosting the IP_3_ pathway. The sensitising effect of Ned-19 (Fig. 5) appears to be specific for ACh-induced Ca²⁺ signalling, as no comparable enhancement was observed in response to GRP stimulation (Fig. 3C). Given that GRP-evoked Ca²⁺ signals are dependent on cADPR, whereas ACh-induced responses involve both the IP₃ and cADPR pathways, the selective potentiation of ACh responses by Ned-19, together with its ability to enhance Ca²⁺ signalling in the presence of the CD38 inhibitor 78c, suggests that the sensitising effect may be mediated through the IP₃-IP₃R signalling axis.

Finally, it is worth remarking on the very different concentrations of both CCK and GRP used in experiments on neurones in the spinal cord and the brain as compared to those employed in our studies of pancreatic acinar cells. Whereas we mostly explored the effects of physiological CCK concentrations in the blood after a meal (1- 10 pM, ref 33) and low concentrations of GRP (125 pM), the concentrations used in neuronal studies were very much higher (typically about 200 nM CCK and 300 nM GRP) (35,36). In the pancreatic acinar cells, such high concentrations would inevitably induce sustained [Ca²⁺]_i_ elevations that would be toxic and initiate processes leading to acute pancreatitis (37,38). Clearly, the pancreatic acinar cells have evolved an extraordinarily high sensitivity to CCK, also demonstrated *in vivo* (71), as well as to GRP. This allows safe Ca²⁺ signaling. Since CCK-elicited IP_3_ production in pancreatic acinar cells only becomes prominent at high un- physiological CCK concentrations (49), it is possible that the requirements for the additional involvement of non-IP_3_ pathways will disappear at the levels of these peptides needed for neuronal activation (35,36). Our result showing that blockage of non-IP_3_ pathways becomes ineffective at high concentrations of CCK and GRP may indicate that this could be the case.

In conclusion, our findings highlight the remarkable complexity and robustness of agonist-evoked signalling in pancreatic acinar cells. Multiple compensatory mechanisms operate in parallel to preserve Ca²⁺ oscillations and, consequently, pancreatic enzyme secretion. This likely reflects the physiological importance of Ca²⁺-driven pancreatic enzyme secretion. Pancreatic digestive enzymes are essential for the breakdown and absorption of nutrients, and failure to produce or secrete these enzymes results in impaired digestion, nutrient acquisition and metabolism (24). The presence of multiple signalling pathways capable of supporting Ca²⁺ oscillations may therefore represent adaptive mechanisms that safeguards exocrine pancreatic function under a range of conditions. On the other hand, the requirement we have demonstrated for activation of more than one intracellular receptor to initiate Ca²⁺ signal generation provides an important protection against unwarranted Ca²⁺ signals and inappropriate enzyme secretion. Most importantly, the additional engagement of non-IP_3_ pathways demonstrated by our experiments allows activation of secretion at very low agonist concentrations. This is important because increasing the concentrations of CCK and GRP towards levels employed in the central nervous system would produce excessive Ca²⁺ signals that have been shown to be toxic (9,10). Avoiding the risk of overstimulation enables safe Ca²⁺ signaling.

## Materials and methods

### Ethical Approval

All animal procedures conducted in this project were carried out in accordance with the Animals (Scientific Procedures) Act 1986 (ASPA, UK) and were approved by the Cardiff University Animal Care and Ethics Committee at Cardiff School of Biosciences. All work complied with institutional and national guidelines governing the ethical use of animals in research.

Wild-type male C57BI/6 mice (aged up to 6 weeks) were obtained from Charles River Laboratories (UK). It is unknown whether our results are valid for female mice, but to avoid potentially confounding variables (hormonal fluctuations, reproductive cycles) we only used male mice. Animals were housed in plastic cages under a controlled 12 h light–dark cycle with corn cob bedding, with ad libitum access to tap water and a standard pelleted chow.

### Materials

Fluo-4/AM was purchased from Thermo Fisher Scientific (Invitrogen, UK). Collagenase V, Cholecystokinin (CCK), Acetylcholine (ACh), Trypsin–chymotrypsin inhibitor and Caffeine were purchased from Sigma-Aldrich. Human Gastrin-releasing peptide (GRP), Ned-19, 78c, and YM-254890 were obtained from Tocris Bioscience (UK). Unless otherwise stated, all other chemicals and reagents were sourced from Sigma-Aldrich.

### Pancreatic Acinar Cell Isolation

Immediately after dissection, as described previously (47), the pancreas was rinsed twice in NaHEPES buffer containing 140 mM NaCl, 4.7 mM KCl, 10 mM HEPES, 1 mM MgCl₂, 10 mM glucose, and 1 mM CaCl₂ (pH 7.2). The tissue was then injected with Collagenase V (28.9 U/mL; Sigma-Aldrich, UK) dissolved in the same buffer and incubated at 37 °C for 7 min in a shaking water bath. Following digestion, the tissue was gently dissociated by repeated pipetting, and the resulting acinar cells were collected and resuspended in fresh NaHEPES buffer. All experiments were performed at room temperature.

### [Ca^2+^]_i_ Measurements

Isolated pancreatic acinar cells were loaded with 5 µM Fluo-4 AM for 50 min at room temperature and subsequently transferred to a perfusion chamber mounted on the stage of a Leica confocal laser-scanning microscope equipped with a 63×, 1.2 NA water-immersion objective. Cells were continuously superfused with either standard NaHEPES solution or Ca²⁺-free NaHEPES solution and stimulated with the indicated secretagogues and pharmacological inhibitors at the concentrations stated. For experiments using CCK or GRP, 0.1% trypsin–chymotrypsin inhibitor was present throughout the recording period. Fluo-4 was excited at 488 nm, and fluorescence emission was collected between 500 and 560 nm at a resolution of 256 × 256 pixels. Images were acquired at a rate of 127 frames per minute. Image acquisition and analysis were performed using Leica Confocal Software (Leica Microsystems, Germany). Changes in intracellular Ca²⁺ concentration are presented as normalised fluorescence ratios (F/F₀).

### Statistical analysis

The intracellular Ca²⁺ signal was quantified as the area under the curve (AUC), calculated over identical recording periods for each secretagogue alone and following co-application of the secretagogue with the indicated pharmacological agent. AUC values were determined using the integration function in GraphPad Prism (version 11.0.2).

Statistical analyses were performed using GraphPad Prism (version 11.0.2). Data distribution was assessed with the Shapiro–Wilk normality test. Normally distributed data were analysed using a two-tailed paired t-test or one-way ANOVA with Tukey’s multiple-comparisons test and are presented as mean ± SEM. Data that did not meet the assumptions of normality were analysed using the Wilcoxon matched-pairs signed-rank test, Kruskal–Wallis test followed by Dunn’s multiple-comparisons test, or Friedman’s test with Dunn’s multiple-comparisons post hoc test, as appropriate. Non-parametric data are presented as median and interquartile range (IQR), unless otherwise stated. Differences were considered statistically significant at P < 0.05.

### Data availability

All data are available upon reasonable request to the authors.

## Author contributions

OHP and MS conceptualized and designed the study and wrote the paper. All experiments were conducted by MS, supervised by JVG and OVG. All data were analysed by MS, OVG and JVG. All authors participated in the revisions of the ms and approved the final version.

## Acknowledgements

MS is a CARA Fellow (Council for At-Risk Academics). OHP was supported by the Medical Research Council (UK) (MR/ J002771/1 and G19/22/2). OVG and JVG were supported by Children with Cancer UK (grants 2017/248 and 2019/288), and Tenovus (PhD2019-23).

